# <u>R</u>AD C<u>apture</u> (Rapture): Flexible and efficient sequence-based genotyping

**DOI:** 10.1101/028837

**Authors:** Omar A. Ali, Sean M. O’Rourke, Stephen J. Amish, Mariah H. Meek, Gordon Luikart, Carson Jeffres, Michael R. Miller

**Author notes:** Corresponding Author: Michael R. Miller Department of Animal Science University of California One Shields Avenue Davis CA 95616.

## Abstract

Massively parallel sequencing has revolutionized many areas of biology but sequencing large amounts of DNA in many individuals is cost-prohibitive and unnecessary for many studies. Genomic complexity reduction techniques such as sequence capture and restriction enzyme-based methods enable the analysis of many more individuals per unit cost. Despite their utility, current complexity reduction methods have limitations, especially when large numbers of individuals are analyzed. Here we develop a much improved restriction site associated DNA (RAD) sequencing protocol and a new method called Rapture (<u>R</u>AD c<u>apture</u>). The new RAD protocol improves versatility by separating RAD tag isolation and sequencing library preparation into two distinct steps. This protocol also recovers more unique (non-clonal) RAD fragments which improves both standard RAD and Rapture analysis. Rapture then uses an in-solution capture of chosen RAD tags to target sequencing reads to desired loci. Rapture combines the benefits of both RAD and sequence capture, i.e. very inexpensive and rapid library preparation for many individuals as well as high specificity in the number and location of genomic loci analyzed. Our results demonstrate that Rapture is a rapid and flexible technology capable of analyzing a very large number of individuals with minimal sequencing and library preparation cost. The methods presented here should improve the efficiency of genetic analysis for many aspects of agricultural, environmental, and biomedical science.

## Introduction

Massively parallel sequencing (MPS) technologies have revolutionized many aspects of agricultural, environmental, and biomedical science (Koboldt *et al.* 2013; Poland and Rife 2012; Shokralla *et al.* 2012; Shendure and Ji 2008). In population biology, MPS enables *de novo* genome assembly for virtually any species (Haussler *et al.* 2009; Alkan *et al.* 2011) and subsequent characterization of within-species genetic variation through whole-genome resequencing (Wheeler *et al.* 2008; Consortium 2010). Although MPS is widely used for whole genome sequencing and resequencing, using MPS to discover and type genetic variation across entire genomes remains prohibitively expensive for many studies (Shendure and Aiden 2012; Sboner *et al.* 2011; Luikart *et al.* 2003).

Because sequencing large amounts of DNA in many individuals can be cost-prohibitive, researchers often interrogate a subset of the genome to reduce the cost per individual (Baird *et al.* 2008). Many genetic studies, such as those characterizing population demography, performing genetic assignment, or describing phylogenetic relationships often require information from a relatively small number of loci (from tens to hundreds). Other studies such as using association mapping to identify loci that influence phenotypic variation or genome scans to describe differential adaptation between populations typically require information from many more loci (from thousands to millions) (Narum *et al.* 2013; Davey *et al.* 2011). Both the number of loci and the number of individuals analyzed contribute to the total cost of genetic analysis. The optimal genetic analysis strategy will vary dramatically by study. Therefore, methods that facilitate flexibility in the number of loci and individuals analyzed are needed for maximizing the efficiency of genetic analysis.

Sequence capture is one method to reduce genome complexity and thereby allow an increased number of individuals to be analyzed with MPS. Genome sequence information is used to design oligonucleotides that facilitate the isolation of desired genomic regions prior to sequencing (Hodges *et al.* 2007; Gnirke *et al.* 2009). Capturing only genomic regions of interest prior to MPS is more economical than sequencing the entire genome for many studies. In-solution capture has facilitated extensive sequencing of target loci across an individual's genome (Gnirke *et al.* 2009). In addition, capture baits designed for one species can often be used in related species due to the conserved nature of functional sequence or a close phylogenetic relationship (Cosart *et al.* 2011). Although capture can generate high sequence depth at targeted loci, the method has drawbacks including a relatively high library preparation cost prior to capture and low multiplexing capacity during capture.

Restriction enzyme-based methods that limit sequencing to a subset of the genome offer an alternative approach to complexity reduction. Examples of restriction site based genomic complexity reduction include restriction site associated DNA (RAD) (Miller *et al.* 2007; Baird *et al.* 2008), reduced representation library sequencing (Van Tassell *et al.* 2008), and genotyping by sequencing (Elshire *et al.* 2011). Different individual restriction enzymes or enzyme combinations can be used to tailor the resolution of complexity reduction. When combined with barcoded adapters, these methods allow large numbers of individuals to be sequenced simultaneously in a single reaction (Baird *et al.* 2008; Hohenlohe *et al.* 2010; Etter *et al.* 2011). Furthermore, the per-individual cost of library preparation can be very low when samples are barcoded and multiplexed early in library construction. Reduced representation sequencing strategies are now being used extensively in conservation, ecological, evolutionary, and agricultural genetic studies (Davey *et al.* 2013; Narum *et al.* 2013; Poland and Rife 2012). However, these methods are much less flexible than sequence capture with respect to controlling the number and location of genomic loci represented after complexity reduction.

Current sequence-based genotyping technologies span a genomic resolution continuum from sequence capture and reduced representation methods to complete genome resequencing. Each technique offers distinct benefits and limitations. Whole genome resequencing provides complete resolution but is cost-prohibitive and unnecessary for many studies involving a large number of individuals. Restriction site based methods offer rapid and inexpensive library preparation for large numbers of individuals but poor flexibility in the number and location of genomic loci analyzed. Sequence capture provides great flexibility with respect to the number and location of genomic loci analyzed but is expensive when applied to large numbers of individuals. New methods that facilitate genotyping of hundreds to thousands of loci in a very large number of individuals would enable many studies that are not currently feasible. Thus, we sought to develop a rapid, flexible, and cost-effective technology that is capable of analyzing a very large number of individuals at hundreds to thousands of loci.

Here we develop a much improved RAD sequencing protocol and a new method called Rapture (<u>R</u>AD c<u>apture</u>). The new RAD protocol improves versatility by separating RAD tag isolation and sequencing library preparation into two distinct steps. This protocol also recovers more unique (non-clonal) RAD fragments which improves both standard RAD and Rapture analysis. Rapture then uses an in-solution capture of chosen RAD tags to target sequencing reads to desired loci. Rapture combines the benefits of both RAD and sequence capture, i.e. very inexpensive and rapid library preparation for many individuals as well as high specificity in the number and location of genomic loci analyzed. Our results demonstrate that Rapture is a rapid and flexible technology capable of analyzing a very large number of individuals with minimal sequencing and library preparation cost. The methods presented here should improve the efficiency of genetic analysis in many areas of biology.

## Results

### A new RAD protocol outperforms the traditional protocol

Our initial goal was to investigate the potential of RAD capture (Rapture) as a flexible and efficient method for sequence-based genotyping. However, our initial Rapture results contained very high PCR duplicate rates (e.g. greater than 90%; data not shown). These could be identified because we used paired-end sequencing and one end of each RAD fragment is generated by a random shearing event (Miller *et al.* 2007; Andrews *et al.* 2013). Upon further investigation, we determined that our RAD libraries contained high clonality even before the capture step (see below). Furthermore, although the traditional RAD protocol (Baird *et al.* 2008; Miller *et al.* 2012) has worked well for us with high-quality DNA samples, the protocol has been inconsistent when using low quality and/or low concentration DNA samples which are frequently encountered in conservation and ecological genetic studies. We reasoned that a new protocol which physically isolates RAD tags from the rest of the genome prior to sequence library preparation would be more robust and reduce clonality (Figure 1A). The new protocol employs biotinylated RAD adaptors that purify the RAD tags after ligation using streptavidin-coated magnetic beads (Miller *et al.* 2007). The purified RAD tags are then used as input to any commercially available library production kit (Figure 1B).

**Figure 1.**
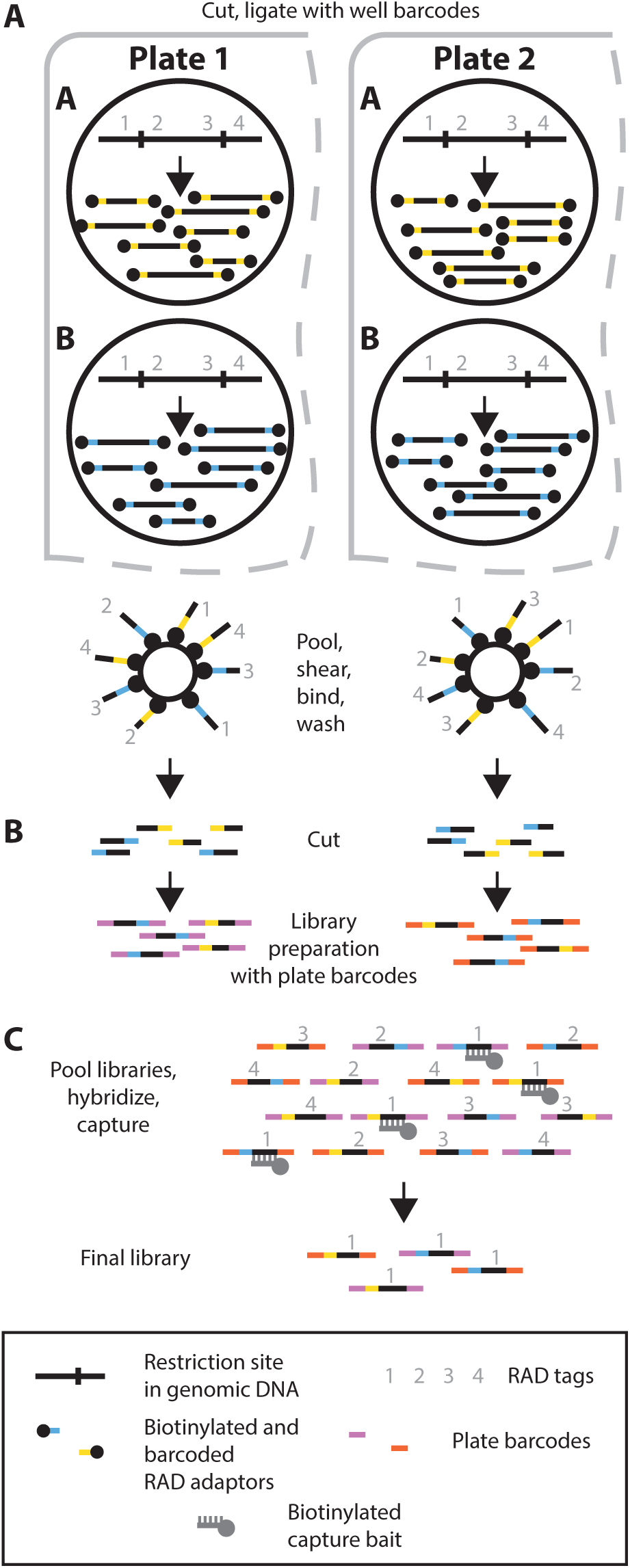
Schematic overview of the new RAD protocol and RAD capture (Rapture) method. (A) RAD tag isolation. Two wells are depicted in each of two different plates. Genomic DNA is digested with a restriction enzyme and ligated to biotinylated well barcode adaptors (yellow and blue bars). (B) RAD tag isolation and library preparation. DNA from each well is pooled platewise, mechanically sheared, and incubated with streptavidin beads. Following washing, DNA is cleaved from the beads leaving the well barcodes. Finally, a library preparation is performed where a unique plate barcode is added (red and purple bars). (C) Rapture. Multiple plate libraries are pooled, hybridized to biotinylated oligonucleotide baits corresponding to the targeted RAD tag loci, and captured to produce the final library enriched for the loci of interest.

To directly compare the new and traditional RAD protocols, we generated and analyzed data from 96 rainbow trout individuals using both procedures. We normalized the sequence data so the analysis of each protocol started with an equal number of reads. We separated the sequence reads according to individual barcode, aligned them, and produced summary statistics to evaluate the new protocol. Specifically, we quantified the average number of sequenced fragments per individual, average number of mapped fragments per individual, and average locus coverage prior to clone removal. The new RAD protocol produced similar numbers of sequenced fragments per individual (means of 1.18 × 10^6^ for the new and 1.24 × 10^6^ for the traditional), and slightly more mapped fragments per individual (9.84 × 10^5^ for the new and 8.06 × 10^5^ for the traditional) (Table 1). In addition, both RAD procedures produced similar distributions of mapped fragments per individual (Figure 2A), but the updated RAD protocol yielded slightly more covered loci per number of sequenced fragments (Figure 2B), as well as better mapping quality of fragments compared to the traditional protocol (Table 1). The DNA used in this experiment was extracted from highly variable fin clips. More consistent samples or more effort in DNA normalization before library preparation could decrease the variance among individuals. These results suggest the updated RAD protocol offers a modest improvement over the traditional RAD protocol even without clone removal.

**Figure 2.**
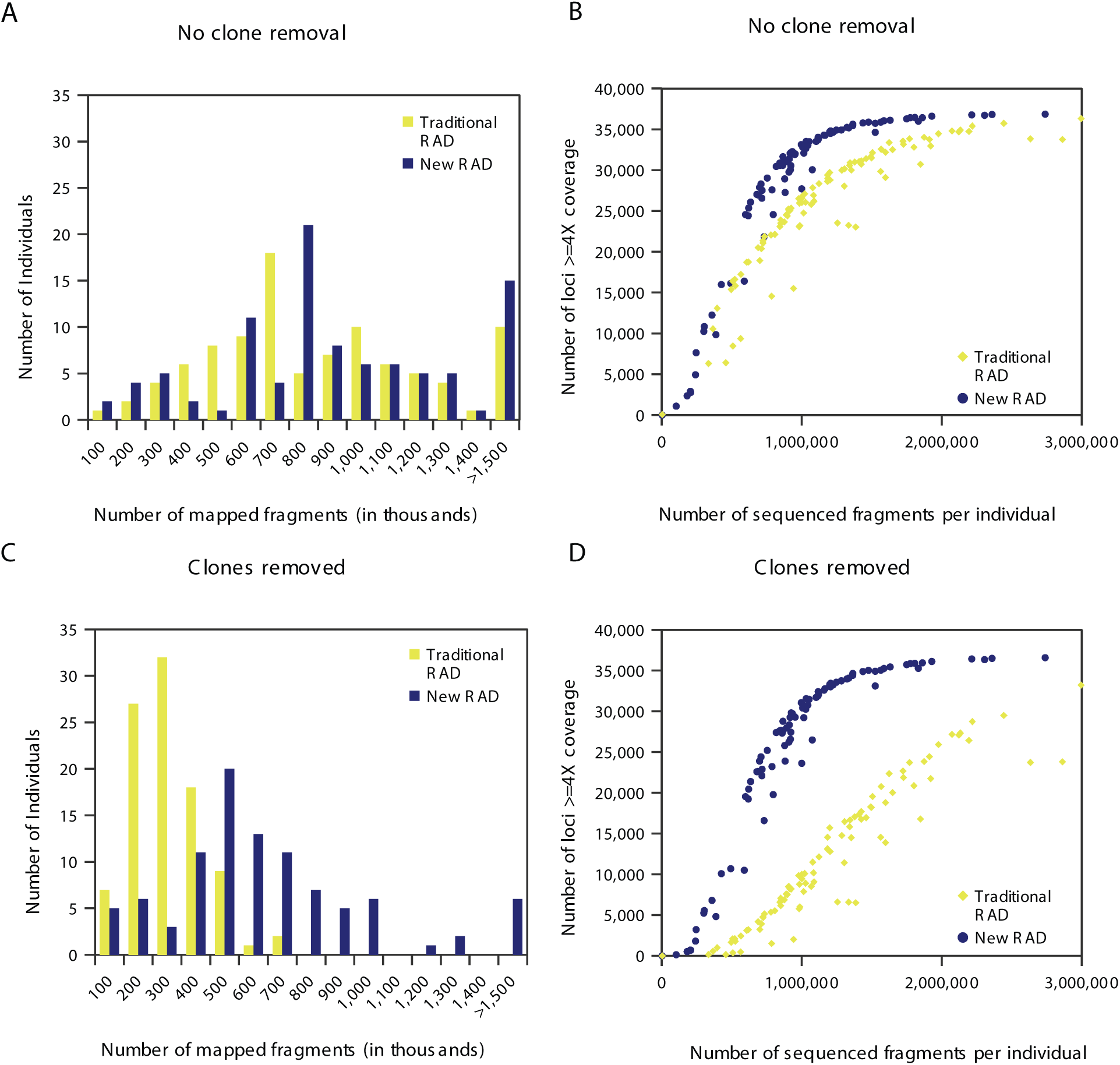
Comparison of RAD sequencing results from traditional and new RAD protocols on 96 individuals. (A) Histogram showing the number of individuals per bin of mapped fragments without clone removal. (B) Scatter plot showing the relationship between number loci covered > = 4X without clone removal and number of sequenced fragments per individual. (C) Histogram showing the number of individuals per bin of mapped fragments with clone removal. (D) Scatter plot showing the relationship between number loci covered > = 4X with clone removal and number of sequenced fragments per individual.

**Table 1.** Comparison of RAD sequencing results from traditional RAD and new RAD protocols applied to the same 96 individual DNA extractions.

|  | Normalized sequenced fragments | Number of Individuals | Average sequenced fragments per individual | Average mapped fragments (no clone removal) | Average locus coverage (no clone removal) | Average mapped fragments (clones removed) | Average locus coverage (clones removed) |
| --- | --- | --- | --- | --- | --- | --- | --- |
| Traditional RAD | 122,356,753 | 96 | 1,236,438 | 805,659 (65.2%*) | 8.23 | 253,381 (20.5%*) | 2.84 |
| New RAD | 122,356,753 | 96 | 1,182,936 | 984,441 (83.2%*) | 9.77 | 624,642 (52.8%*) | 7.03 |

\*Percentage of average sequenced fragments per individual

To determine if the updated RAD protocol produces fewer PCR duplicates, we removed clonal sequences prior to determining alignment statistics and locus coverage. Strikingly, the traditional RAD protocol produced high numbers of clones which substantially reduced the number of unique mapped fragments per individual (Figure 2C). With clone removal, the average locus coverage in the traditional protocol was reduced to 2.84X (a 65% loss of coverage) whereas in the new protocol coverage was 7.03X (a 28% loss) (Table 1). Finally, the number of fragments required for the traditional protocol to reach similar coverage levels of the updated protocol is substantially higher (Figure 2D). Our new RAD protocol significantly improved the average number of mapped fragments, the coverage per locus, and the number of loci covered per barcoded fragment. We conclude that the new RAD protocol offers substantial improvements over the traditional protocol.

### Rapture produces high coverage from minimal reads per individual

To test the Rapture method, we designed and synthesized 500 RNA baits complementary to specific rainbow trout RAD tags distributed across the 29 chromosomes and performed RAD capture. We produced RAD sequencing libraries for each of three 96-well plates using the new protocol. Each individual within a plate had a unique well barcode and each plate had a unique plate barcode. This allowed the three RAD libraries to be combined into a single library containing a total of 288 individuals. We then performed a single capture reaction with the recommended bait concentration on the combined library (Figure 1C). We sequenced both pre-and post-capture versions of the combined library in a small fraction of an Illumina lane (10%) which produced approximately 20 million sequenced fragments for each of the pre-and post-capture libraries. This experimental design provides a direct comparison between RAD and Rapture and simulates the sequencing of thousands of individuals in an entire single lane.

To evaluate the Rapture results, we performed alignments, clone removal, and generated a number of summary statistics. As above, we quantified the average sequenced fragments per individual, the average mapped fragments per individual, and the average locus coverage. The distribution of sequence reads per individual were consistent between pre-capture (RAD) and Rapture (Figure 3A). However, the average coverage at the captured loci was remarkably different between the RAD and Rapture protocols: RAD produced a 0.43X average coverage whereas Rapture produced 16X average coverage (Table 2). Strikingly, the number of sequenced fragments per individual required for 4X or greater coverage of the captured loci is approximately 30,000 for Rapture, while pre-capture (RAD) failed to cover the captured loci to any extent (Figure 3B). Finally, as expected, the Rapture procedure did not significantly enrich for non-targeted RAD loci (Figure 3C). We conclude that Rapture effectively targets specific RAD loci for high-coverage with minimal sequencing.

**Figure 3.**
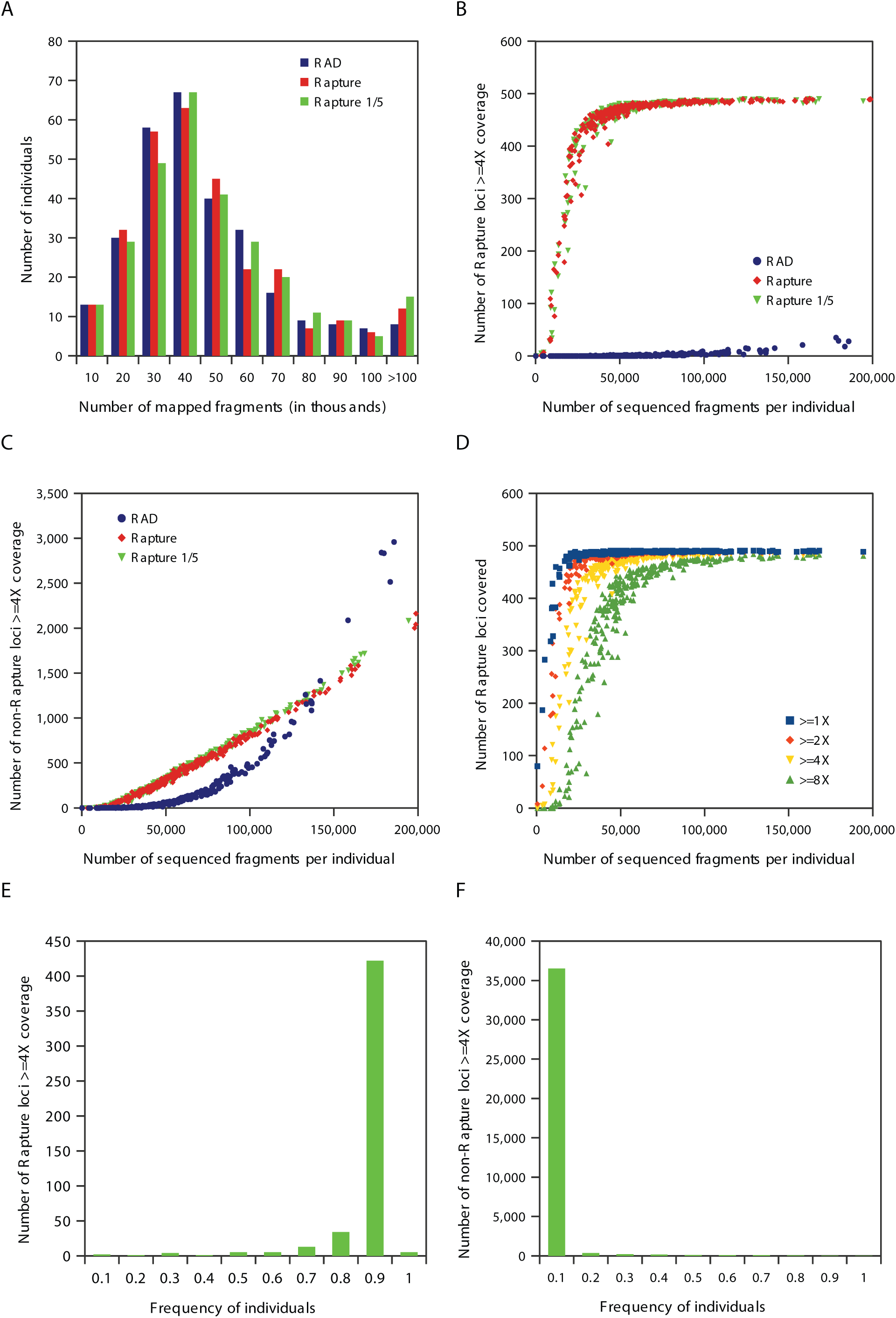
Comparisons of RAD, Rapture, and Rapture with one-fifth bait concentration (Rapture 1/5) sequencing results with clone removal on 288 individuals. (A) Histogram showing the number of individuals per bin of mapped fragments. (B) Scatter plot showing the relationship between number Rapture loci covered > = 4X and number of sequenced fragments per individual. (C) Scatter plot showing the relationship between number non-Rapture loci covered > = 4X and number of sequenced fragments per individual. (D) Scatter plot showing the relationship between number Rapture loci covered at select levels and number of sequenced fragments per individual for Rapture 1/5. (E) Histogram showing number of Rapture loci covered > = 4X per bin individuals for Rapture 1/5. (F) Histogram showing number of non-Rapture loci covered > = 4X per bin individuals for Rapture 1/5.

**Table 2.** Comparison of new RAD, Rapture, and Rapture with one-fifth bait concentration (Rapture 1/5) sequencing results.

|  | Normalized sequenced fragments | Number of individuals | Average sequenced fragments per individual | Average mapped fragments (no clone removal) | Average mapped fragments (clones removed) | Average non-Rapture locus coverage (clones removed) | Average Rapture locus coverage (clones removed) |
| --- | --- | --- | --- | --- | --- | --- | --- |
| RAD | 21,879,887 | 288 | 57,630 | 46,655 (81.0%*) | 41,820 (72.6%*) | 0.45 | 0.43 |
| Rapture | 21,879,887 | 288 | 65,978 | 56,555 (85.7%*) | 42,288 (64.1%*) | 0.40 | 16.01 |
| Rapture 1/5 | 21,879,887 | 288 | 66,540 | 58,075 (87.3%*) | 44,380 (66.7%*) | 0.42 | 16.38 |

\*Percentage of average sequenced fragments per individual

To simulate performing Rapture with an increased number of individuals in a single capture reaction, we performed a second capture with the same libraries as above but used only one-fifth the recommended concentration of capture baits. The RAD capture performed with a one-fifth capture bait concentration behaved identically to the original Rapture data (Figure 3A, 3B, and 3C). The amount of sequencing needed to gain specific coverage levels is shown in Figure 3D. A high percentage of Rapture loci are covered at 4X or greater for more than 90% of the individuals, whereas non-Rapture loci were covered at 4X or greater in less than 10% of the individuals (Figures 3E and 3F). With both Rapture trials producing identical results, we conclude that Rapture can process a minimum of 500 loci from 1440 individuals (5 × 288) per capture reaction.

### Rapture reveals population structure in Fall River rainbow trout

We next investigated the suitability of Rapture sequence data for genetic analysis by discovering and genotyping SNPs using clone-removed alignments from the Rapture one-fifth bait concentration experiment described above. We first determined the distance from the restriction site for each SNP discovered in the Rapture loci. Sequencing was done with 100 bp reads but the first 16 bases on the cut site end of the sequenced fragment were removed because they contained the barcode and partial cut site. Additionally, the shearing and size selection protocol used for these experiments produced fragments up to 500 bases in length. Therefore, positions beyond 84 bases from the cut site should have lower coverage. As expected, most SNPs were discovered near the cut site due to higher sequencing depth generated on that end of the DNA molecule (Figure 4A). We discovered 637 SNPs within the first 84 bases following the cut site and 1507 SNPs between bases 85-500 (Table 3). The exact shearing, size selection, and sequencing parameters could be adjusted in future experiments to influence the number of discovered SNPs.

**Figure 4.**
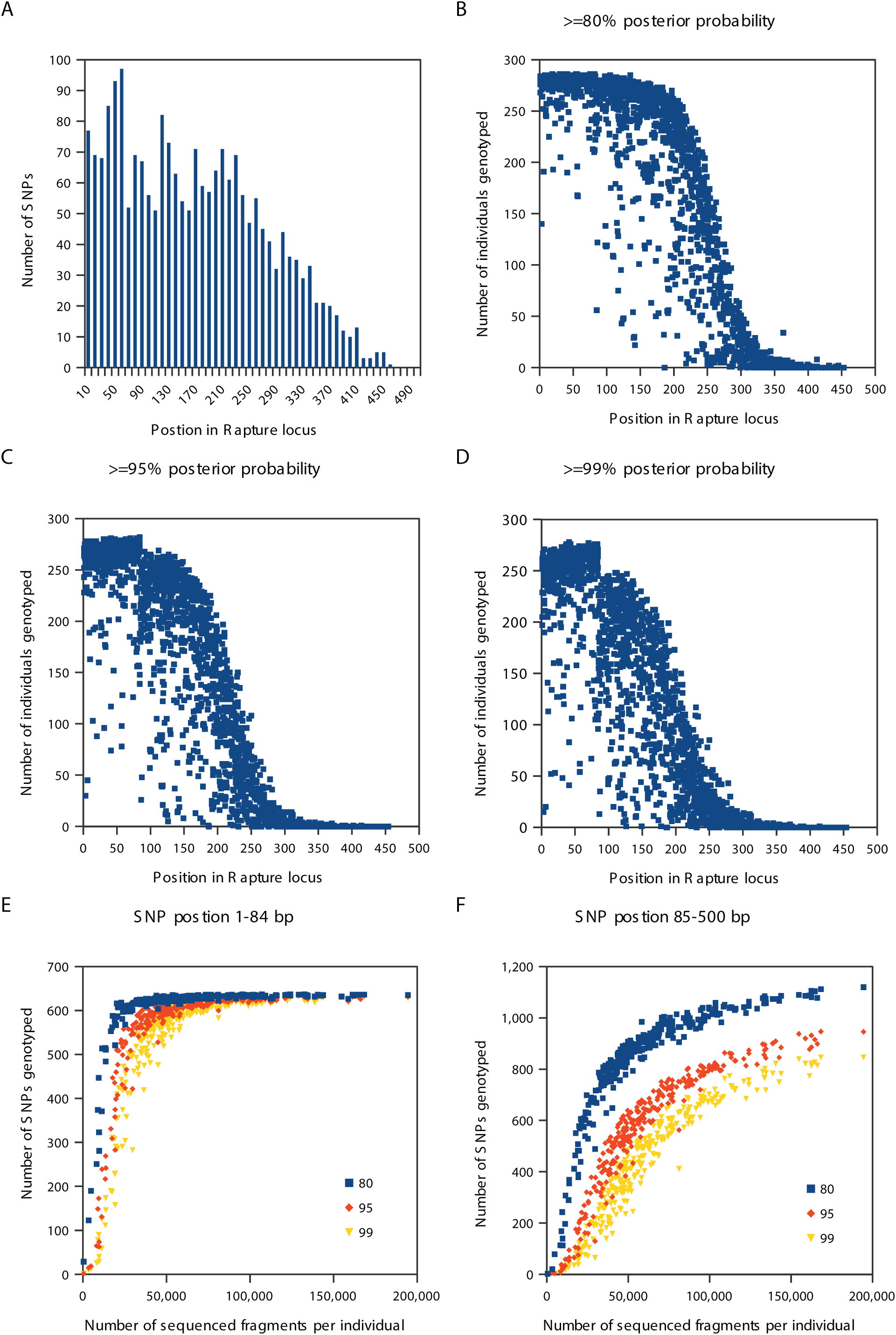
SNP discovery using Rapture with one-fifth bait concentration data. (A) Histogram showing the number of SNPs per bin of position in Rapture locus. (B-D) Scatter plots showing the relationship between the number of individuals genotyped and SNP position using different posterior probability cutoffs. (E) Scatter plot showing the relationship between the number of SNPs genotyped and number of sequenced fragments per individual for SNPs in position 1–84. (F) Scatter plot showing the relationship between the number of SNPs genotyped and number of sequenced fragments per individual for SNPs in position 85–500.

**Table 3.**
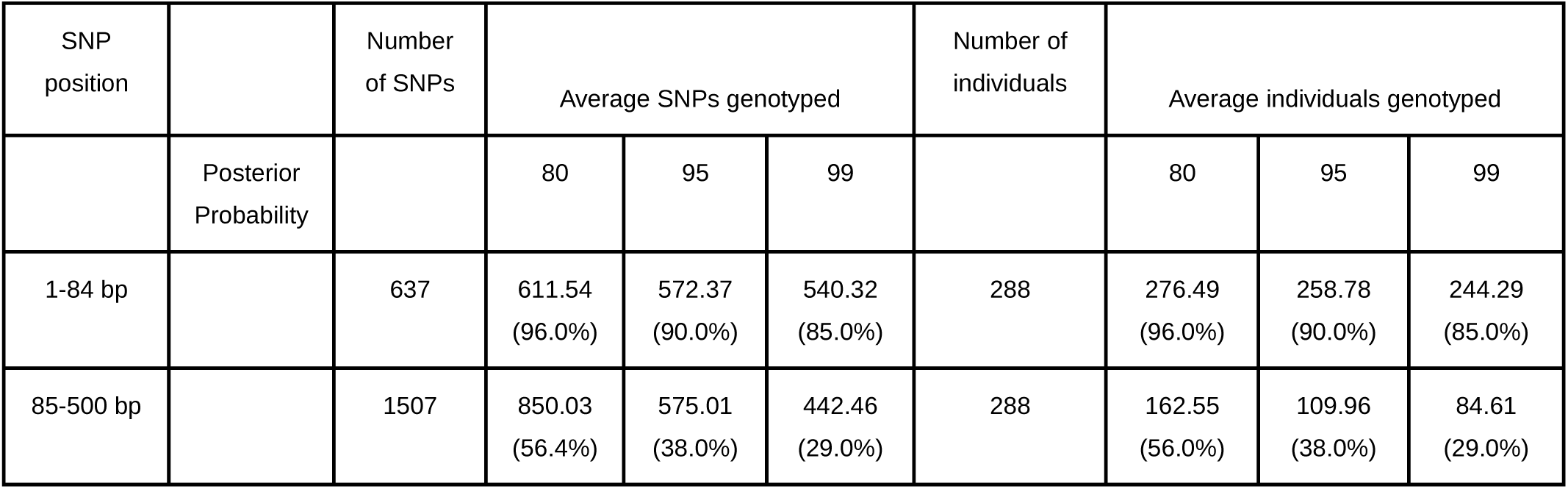
Comparison of genotyping rate and position of SNP in Rapture locus.

| SNP position |  | Number of SNPs | Average SNPs genotyped |  |  | Number of individuals | Average individuals genotyped |  |  |
| --- | --- | --- | --- | --- | --- | --- | --- | --- | --- |
|  | Posterior Probability |  | 80 | 95 | 99 |  | 80 | 95 | 99 |
| 1-84 bp |  | 637 | 611.54<br>(96.0%) | 572.37<br>(90.0%) | 540.32<br>(85.0%) | 288 | 276.49<br>(96.0%) | 258.78<br>(90.0%) | 244.29<br>(85.0%) |
| 85-500 bp |  | 1507 | 850.03<br>(56.4%) | 575.01<br>(38.0%) | 442.46<br>(29.0%) | 288 | 162.55<br>(56.0%) | 109.96<br>(38.0%) | 84.61<br>(29.0%) |

We then plotted the number of successfully genotyped individuals for each SNP along the length of the RAD fragments using different genotype posterior probability cut-offs. We found that SNPs within the first 84 bases were successfully genotyped at each cut-off level used (Figure 4B, 4C, and 4D). Also, the number of SNPs genotyped within the first 84 bases given a minimal number of reads (approximately 25,000) for each individual is extremely high (Figure 4E). SNPs located after the first 84 bases required more sequencing to approach saturation in the number of individuals with called genotypes (Figure 4F). We conclude that Rapture facilitates versatile and high quality SNP discovery and genotyping.

To test the utility of Rapture-generated genotypes for investigating population structure, we calculated a covariance matrix from 273 well-genotyped individuals and performed a principal component analysis (Figure 5A and 5B, see Materials and Methods). The wild rainbow trout used in the study originated from the Fall River watershed in Shasta County, California. Fin clips were collected from adult fish with unknown birth locations in the Fall River system as well as from juvenile fish that were born at known upstream spawning locations (Figure 5C). Strikingly, the first principal component separated two distinct groups corresponding to individuals born in Bear Creek and the spring-fed spawning locations (Thousands Springs and Spring Creek) (Figure 5A). Furthermore, genetic differentiation is even apparent between spring-fed spawning sites as individuals from Thousand Springs and Spring Creek separate on the third component (Figure 5B). Thus, Rapture facilitated the discovery of population structure on a fine spatial scale within a closed watershed. We conclude that Rapture is a useful tool for characterizing population structure.

**Figure 5.**
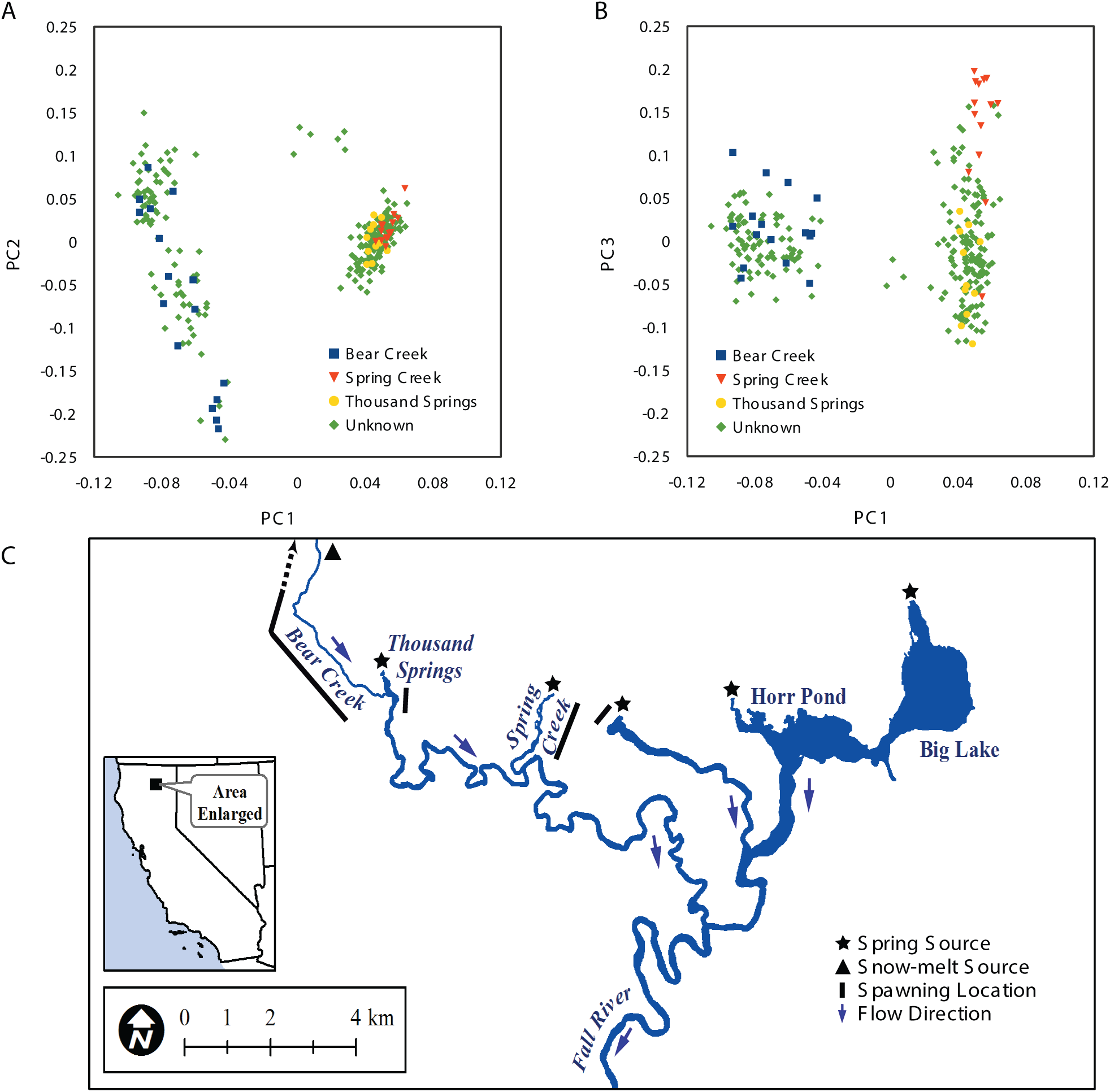
Principal component analysis of Rapture genotyping results from Fall River rainbow trout. Individuals labeled based on birth location. Individuals with known birth locations were collected as juveniles near spawning grounds. Other individuals were collected as adults throughout the system below the spawning grounds. (A) Scatter plot showing the first two principal components. (B) Scatter plot showing the first and third principal components. (C) Fall River map.

## Discussion

### A new RAD protocol is superior to the traditional protocol

PCR clones are a serious problem in sequence-based genotyping because they produce incorrect genotype calls (Andrews *et al.* 2014; Hohenlohe *et al.* 2013). Clones are easily detected with some RAD sequencing protocols due to a random shearing step that produces a unique breakpoint in the DNA fragment. Because the Rapture protocol used here relied on two PCR amplification steps (one during the RAD library construction, and another subsequent to the capture), clonality present in the initial RAD libraries is exacerbated in the final post-capture library. Thus, we sought to minimize the level of clonality as much as possible in the RAD libraries. One way to do this is to simply use more genomic DNA, however our samples are often finite and yield low DNA concentrations due to small sized or degraded tissue. Therefore, we developed an improved RAD sequencing procedure to maximize RAD tag diversity.

Our redesigned RAD protocol employs physical enrichment of RAD tags rather than PCR-based enrichment. The new RAD protocol outperforms the traditional protocol by yielding increased numbers of mapped fragments, better coverage per locus, and requires less sequence data to achieve the same coverage. The physical separation of RAD tags from other genomic fragments captures more unique (non-clonal) RAD fragments than the older method of PCR enrichment. We have now used the new RAD protocol on many diverse samples. The new protocol consistently produces higher concentration libraries using the same input DNA and PCR cycles. Furthermore, the new protocol is much more robust. For example, with low concentration and/or low quality samples, failed libraries were fairly common when using the old protocol but are virtually non-existent with the new protocol. A possible explanation for the relatively poor performance of the traditional protocol is that the PCR template contains a very high percentage of non-amplifiable DNA fragments that have divergent “Y” adapters on both ends.

The separation of RAD tag isolation and sequencing library preparation into two distinct steps offers a significant benefit in addition to reduced clonality. New advancements in sequencing library preparation reagents (such as hairpin loop adapters) were incompatible with the traditional RAD protocol due the integrated steps of RAD tag isolation and library preparation. However, the updated RAD protocol produces barcoded, double-stranded, sheared DNA that can be used as input material for any library preparation protocol on any sequencing platform that accepts fragmented DNA. This new protocol should also be compatible with PCR-free library preparation kits which would completely remove PCR duplicates. In conclusion, the new physical RAD tag isolation procedure generates higher quality data, is more cost-effective, and allows more flexibility for library production than the traditional protocol.

### Rapture combines the benefits of RAD and sequence capture

RAD sequencing excels at sample multiplexing during library preparation because each plate of 96 samples is rapidly combined into a singe tube after barcoding and processed as a single reaction. Furthermore, the library from each plate also receives a unique plate barcode which allows processing multiple plate libraries in a single capture reaction. Thus, the procedure we present is scalable to many thousands of samples. The in-solution capture step in Rapture uses a commercially available kit to selectively isolate the desired RAD loci for sequencing. Here we targeted 500 loci but any number of capture baits could be used to target more or fewer RAD loci. The MYcroarray commercial product we used offers up to 200,000 unique baits per kit.

Our results demonstrate the potential for Rapture to genotype thousands of individuals in a single sequencing reaction. We analyzed Rapture data obtained from approximately 10% of one Illumina HiSeq lane (approximately 20 million reads) and still achieved over 16X coverage across 500 Rapture loci in 288 individuals. Therefore, sequencing 500 loci at approximately 4X, 8X and 16X coverage could be achieved by multiplexing 11,520, 5,760, and 2,880 individuals per lane of sequencing, respectively. Recent improvements in sequence outputs with new Illumina machines will allow even higher levels of multiplexing. Furthermore, even coverage as low as 2X can provide sufficient data for many questions when using probabilistic genotyping approaches (Nielsen *et al.* 2011).

### Experimental design considerations for Rapture

Bait design for Rapture can be obtained from a reference genome, prior RAD data, or by RAD sequencing a subsample of individuals to be used for Rapture. Once candidate loci are identified, some number of loci are chosen to design a custom bait library kit used for sequence capture. This number is based on the aims and budget of the study. RAD libraries could be generated and baits designed adjacent to 8 bp (such as SbfI) or 6 bp (such as PstI) restriction sites. Either way, RAD tags can be chosen to provide a random representation of the genome or designed with specific requirements depending on experimental needs. Requirements for RAD tags can be based on molecular constraints and/or genetic information from prior analysis such as linkage with respect to other RAD tags, linkage with respect to phenotype, polymorphism, paralogy, position in the genome or genetic maps (e.g. near genes), etc.

If these factors are not considered, the quality and quantity of Rapture sequence data may be diminished and potentially insufficient to answer the biological questions of interest. Several other factors could produce low quality data such as the molecular biology of sequence capture (sub-optimal bait design), the designing of Rapture baits that have paralogous (or highly similar) sequences represented throughout the genome (off-target capture and sequencing), and the total number of RAD loci chosen or individuals sequenced (an inappropriate relationship between the numbers of individuals, the numbers of loci, and amount of sequencing). Therefore, Rapture loci discovery and bait design is an important first step for a successful experiment.

### Genetic population structure of Fall River rainbow trout

We demonstrated the successful use of the new RAD protocol and Rapture by detecting genetic population structure within a small geographic area (the Fall River watershed of Northern California) using a relatively small number of sequenced fragments per individual. We used Rapture to generate sequence at 500 RAD loci, thoroughly interrogating over 40,000 bases per individual. By knowing the hatching location of juvenile fish, we could infer origin of adult fish to Bear Creek or the spring spawning locations. We were very surprised to discover significant population structure in the rainbow trout from such a small watershed.

The Fall River flows approximately 34 kilometers from the uppermost spring and consists of two distinct sources of water: many individual spring inputs and a single rain/snow-melt stream (Figure 5C). A dam just upstream of the Pit River confluence blocks upstream fish passage, so the Fall River trout population is self-sustaining and does not receive migrants from outside locations. Bear Creek is an ephemeral tributary of the Fall River that fluctuates from zero to approximately 28 cubic meters per second (cms) during the year, depending on precipitation events and snowpack. The major water source for the Fall River are the multiple springs that discharge at a relatively constant rate of approximately 35 cms. The large contribution of water from springs keeps the Fall River within a constant temperature range and flow regime throughout the year, with the exception of high flow events from Bear Creek. The ephemeral and perennial differences between the spawning locations are likely responsible for producing the genetic differentiation between the two major distinct groups discovered here.

## Materials and Methods

### Genomic DNA extractions

We developed an economical high throughput DNA extraction method using Agencourt Ampure XP beads and a Liquidator 96 Manual 96-well Pipettor (Rainin). Liftons buffer (80 µl, 100 mM EDTA, 25 mM tris-HCl pH 7.5, 1% SDS) was added to each well of a 96 well plate. Fin clips measuring 2-25 mm^2^ were placed into each well. Liftons buffer (40 µl) containing 0.075 M DTT and 4.2 mg/ml Proteinase K was added to each well. After mixing, the plate was sealed and incubated at 55ºC overnight to generate a crude DNA lysate. To a new plate containing 45 µl Hybridization buffer (2.5 M NaCl, 20% PEG 8000, 0.025 M DTT) and 15 µl Agencourt AMPure XP beads (Beckman Coµlter, A63881), 45 µl of the crude lysate was added. After thorough mixing, the plate was incubated for 5 min and then placed on a magnet. The supernatant was aspirated and discarded. The plate was removed from the magnet and 150 µl freshly prepared 80% ethanol was used to resuspend the Ampure beads. Two additional 80% ethanol washes were performed. The beads were allowed to air dry while on the magnet and then a volume (20-100 µl) of low TE (10 mM tris-HCl pH 7.5, 0.1 mM EDTA) was used to elute the DNA from the beads.

### Traditional RAD

For each sample, genomic DNA (50 ng) was digested with 2.4 units of SbfI-HF (New England Biolabs, NEB, R3642L) at 37ºC for 1 h in a 12 µl reaction volume buffered with 1X NEBuffer 4 (NEB, B7004S). After heat inactivation at 80ºC for 20 minutes, 2 µl indexed SbfI/PstI P1 RAD adapter (10 nM) was added to each sample (see Supplemental File 1 for sequences). To ligate the adaptors to the cleaved genomic DNA, 2 µl of ligation mix (1.28 µl water, 0.4 µl NEBuffer 4, 0.16 µl rATP (100 mM, Fermentas R0441), 0.16 µl T4 DNA Ligase (NEB, M0202M) was added. Ligations were performed at 20ºC for 1 h followed by incubation at 65ºC for 15 min to inactivate the ligase. For each of the 96 samples, 5 µl was pooled and precipitated with 1X Agencourt AMPure XP beads (Beckman Coµlter, A63881). The remaining sample was reserved for additional library preparation if desired. The pooled DNA was resuspended in 210 µl low TE and sheared in a Bioruptor NGS sonicator (Diagenode). We used 9 cycles of 30 sec on/90 sec off and evaluated the shearing efficiency with a fragment analyzer (Advanced Analytical Technologies). Additional shearing cycles were performed as necessary. The sheared DNA was then concentrated to 55.5 µl using Ampure XP beads.

The concentrated DNA was used as template in the NEBNext Ultra DNA Library Prep Kit for Illumina (NEB E7370L; version 1.2) with following modifications. Instead of using the supplied Illumina adaptor, we ligated a custom P2 adaptor onto the fragments. The indexed P2 was prepared by annealing an NEBNext Mµltiplex Oligo for Illumina (NEB, E7335L) to the oligo GATCGGAAGAGCACACGTCTGAACTCCAGTCACIIIIIIATCAGAACA*A (the * represents a Phosphorothioated DNA base). We omitted the USER enzyme step and used a universal P1 RAD primer (AATGATACGGCGACCACCGAGATCTACACTCTTTCCCTACACGAC*G) and a universal P2 RAD primer (CAAGCAGAAGACGGCATACG*A) instead of the included NEBNext oligos for the final amplification.

### New RAD Protocol

For each sample, genomic DNA (50 ng) was digested with 2.4 units of SbfI-HF at 37ºC for 1 h in a 12 µl reaction volume buffered with 1X NEBuffer 4 (Note: More DNA can and should be used when available. We have successfully used 200 ng per sample with this exact protocol.). After heat inactivation at 80ºC for 20 minutes, 2 µl indexed SbfI/PstI biotinylated RAD adapter (50 nM) was added to each sample (see Supplemental File 1 for sequences). The new RAD adapters feature 8 bp hamming barcodes (Kozarewa and Turner 2011). To ligate the adaptors to the cleaved genomic DNA, 2 µl of ligation mix (1.28 µl water, 0.4 µl NEBuffer 4, 0.16 µl rATP, 0.16 µl T4 DNA Ligase) was added. Ligations were performed at 20ºC for 1 h followed by incubation at 65ºC for 15 min to inactivate the ligase. For each of the 96 samples, 5 µl was pooled and precipitated with 1X AMPure XP beads. The remaining sample was reserved for additional library preparation if desired. The pooled DNA was resuspended in 210 µl low TE and sheared in a Bioruptor NGS sonicator. We used 9 cycles of 30 sec on/90 sec off and evaluated the shearing efficiency with a fragment analyzer. Additional shearing cycles were performed as necessary.

We used Dynabeads M-280 streptavidin magnetic beads (Life Technologies, 11205D) to physically isolate the RAD-tagged DNA fragments. A 30 µl aliquot of Dynabeads was washed twice with 100 µl of 2X binding and wash buffer (10 mM Tris-HCl (pH 7.5), 1 mM EDTA pH 8.0, 2 M NaCl). The Dynabeads were resuspended in a volume of 2X binding and wash buffer equivalent to the sheared DNA volume from above. The bead/DNA mixture was incubated at room temperature for 20 min with occasional mixing. The beads were washed twice by placing the tube on a magnetic rack, removing the supernatant and resuspending the beads in 150 µl 1X binding and wash buffer (5 mM Tris-HCl (pH 7.5), 0.5 mM EDTA pH 8.0, 1 M NaCl). Two additional washes were performed using 56ºC 1X binding and wash buffer. Two additional washes were performed using 1X NEBuffer 4. The beads were resuspended in 40 µl 1X NEBuffer 4 containing 2 µl SbfI-HF. After incubation at 37ºC for one hour, the supernatant containing the liberated DNA was removed and precipitated with 1X AMPure XP beads. The DNA was eluted in 55.5 µl of low TE and used in NEBNext Ultra DNA Library Prep Kit for Illumina with no modifications.

### Sequence capture of RAD tags for Rapture

Baits were designed based on sequence from a previous experiment that identified 40,649 high quality SbfI RAD loci in rainbow trout (Miller *et al.* 2012). A list of potential baits were chosen based on optimal GC content and minimal sequence similarity to other loci. From that list, 500 loci were chosen for bait design such that all linkage groups (Miller *et al.* 2012) had approximately equal coverage. We then ordered baits from MYcroarray and used the MYbaits protocol supplied with the capture probes. The only modification we made was to use universal primers in the final library amplification because we combined several libraries, each made with unique barcoded primers. The universal primers had the sequence: AATGATACGGCGACCACCGAGATCTACACTCTTTCCCTACACGAC*G and CAAGCAGAAGACGGCATACG*A (the * represents a Phosphorothioated DNA base).

### Sequencing, alignments, and coverage analysis

Libraries were sequenced with paired-end 100 bp reads on an Illumina HiSeq 2500. For each analysis, the sequencing libraries were randomly subsampled to produce an equivalent number of reads for each library. The libraries were demultiplexed by requiring reads to have a perfect barcode match as well as a perfect partial restriction site match for assignment to an individual. To demultiplex the traditional RAD data, only the first reads were searched for a barcode and partial restriction site. In the new RAD protocol, the barcode and partial restriction site can be on either read. Therefore, both reads were searched during demultiplexing. Rare cases in which a barcode and partial restriction site were present on both reads were discarded.

Reads were aligned to the rainbow trout reference genome assembly (Berthelot *et al.* 2014) using the `aln` algorithm implemented in BWA (Li and Durbin 2009). Of the 40,649 previously discovered RAD loci (Miller *et al.* 2012), 38,144 were present and represented only once in the reference genome. 496 of the 500 Rapture baits were present and represented only once in the reference genome. Therefore, the RAD analyses examined 38,144 loci and the Rapture analyses examined 496 loci.

Coverage statistics were obtained by analyzing these loci with Samtools (Li et al. 2009). The alignments were filtered for proper pairs with Samtools `view` and PCR duplicates were removed with Samtools `rmdup,` except in first analysis comparing the new and traditional RAD protocols. Samtools `flagstat` was used to determine the number of fragments, number of alignments, and number of unique alignments (clones removed). Samtools `depth` was used to determine coverage per locus.

### SNP discovery and population genetic analysis

Filtered BAM files generated from the above analysis were used in ANGSD (Korneliussen *et al.* 2014) for SNP discovery, genotype posterior calculation, and population genetic analysis. SNP discovery was conducted on sites with a minimum base quality of 20 and a minimum mapping quality of 20. Sites were characterized by estimating their minor allele frequency using a uniform prior and the Samtools genotype likelihood model (Li 2011). Sites were designated polymorphic if the SNP p-value was less than or equal to 1e-6. Individuals were genotyped at each site using posterior probability cut-offs of 0.80, 0.95, and 0.99. We then parsed the output files to determine the position of each SNP relative to the RAD restriction site and the numbers of individuals genotyped for each SNP.

To perform the principal component analysis, we subsampled the BAM files of each individual to the same number of mapped fragments (10,000), which left 273 of the original 288 individuals. An ANGSD genotype posterior output was generated with a uniform prior and filtered by mapping quality (20), base quality (20), SNP p-value (1e-6), minimum minor allele frequency (0.05), and minimum individuals (220). This called genotype output was used to calculate a covariance matrix with ngsTools `ngsCovar` (Fumagalli *et al.* 2013). Principal component axes summarizing population genetic structure were derived from this co-variance matrix by eigenvalue decomposition.

## Supporting Information

Supplemental File 1. RAD protocol adapter sequences.

Supplemental File 2. Bait sequences.

Supplemental File 3. Rainbow trout sample information.

## Acknowledgments

Fall River Conservancy in particular Andrew Braugh; California Trout; S.D. Bechtel, Jr. Foundation; 1000 Springs Ranch; Steve McCanne; California Department of Fish and Wildlife-Heritage and Wild Trout Program; Members of the Genetic Diversity Research Group and UC Davis Watershed Science Center in particular Eric Holmes, Daniel Prince, and Ismail Saglam for help with sample collection and data analysis; Iwanka Kozarewa for hamming barcode sequences. Gordon Luikart and Stephen Amish were supported by grants from National Science Foundation (DEB-1258203) and Montana Fish Wildlife and Parks. This work used the Vincent J. Coates Genomics Sequencing Laboratory at UC Berkeley, supported by NIH S10 Instrumentation Grants S10RR029668 and S10RR027303.

